# Plasma Contains Ultra-short Single-stranded DNA in Addition to Nucleosomal cfDNA

**DOI:** 10.1101/2021.08.22.457243

**Authors:** Jordan Cheng, Marco Morselli, Wei-Lun Huang, You Jeong Heo, Thalyta Pinheiro-Ferreira, Feng Li, Fang Wei, David Chia, Hua Jun, Kenneth Cole, Wu-Chou Su, Matteo Pellegrini, David TW Wong

**Affiliations:** School of Dentistry, University of California, Los Angeles, Los Angeles; Department of Molecular, Cellular & Developmental Biology, David Geffen School of Medicine, University of California, Los Angeles, Los Angeles; Center of Applied Nanomedicine, National Cheng Kung University, Tainan, Taiwan; The Samsung Advanced Institute for Health Sciences & Technology (SAIHST), Samsung Medical Center, Sungkyunkwan University School of Medicine, Seoul, Republic of Korea; Department of Pathology, David Geffen School of Medicine, University of California, Los Angeles, Los Angeles; National Institute of Standards and Technology, Gaithersburg, Maryland

**Author notes:** Joint-first author - These authors contributed equally to this work (Jordan Cheng, Marco Morselli).

## Abstract

Plasma cell-free DNA is a widely used biomarker for diagnostic screening. We introduce uscfDNA-seq, a single-stranded cell-free DNA NGS pipeline, which bypasses previous limitations to reveal a novel population of ultrashort single-stranded cell-free DNA in plasma with a modal size of 50nt. This species of cfDNA aligns predominantly to the nuclear genome and could potentially be used for novel biomarker discovery.

## MAIN TEXT

In liquid biopsy, cell-free DNA (cfDNA) analysis has typically been focused on the mono-nucleosomal cfDNA (mncfDNA) biomarker of ∼160bp in length. However, the current impression of cfDNA is influenced by biases in nucleic acid extraction and library preparation. The development of single-stranded library preparation methods suggests that in addition to mncfDNA, there are shorter cfDNA fragments (<100bp), which are either single-stranded or nicked dsDNA in plasma.^1,2^ Previous studies indicate that size selecting shorter fragments of cfDNA can be used to enrich for mutant-containing cfDNA fragments in late-stage cancer patients.^3^ Next-generation sequencing approaches examining differences in plasma cfDNA fragment lengths have revealed distinct fragment-profiles in cancer patients.^4^ Additionally, various groups have examined cfDNA strandedness as a diagnostic indicator.^5,6^ With these considerations, ultrashort single-stranded cell-free DNA (uscfDNA) is an unexamined cfDNA entity with clinical potential. Previously, nucleic acid extraction kits were not designed to efficiently retain low-molecular cfDNA (<100bp) regardless of strandedness.^7^ Thus, an effective ultrashort ssDNA cfDNA extraction method which retains low-molecular ultrashort-cfDNA coupled with single-stranded library preparation could reveal more about cfDNA population in the 25-100bp region.

To address this, we introduce an ultrashort single-stranded cfDNA sequencing pipeline (**uscfDNA-seq**) **(Fig 1A and B)**. This pipeline incorporates an ultrashort single-stranded cfDNA **(uscfDNA)** optimized extraction method and single-stranded library preparation. The extraction method utilizes both Solid Phase Reversible Immobilization magnetic beads (SPRI) and phenol:chloroform:isoamyl alcohol to retain low molecular weight fragments in plasma. It leverages a high ratio of isopropanol which generates a DNA-phobic environment to precipitate nucleic acids and proteins before isolating the aqueous nucleic acid-containing portion with phenol:chloroform isoamyl alcohol. Subsequent magnetic bead washes help retain the uscfDNA and remove unwanted constituents.

**Figure 1.**
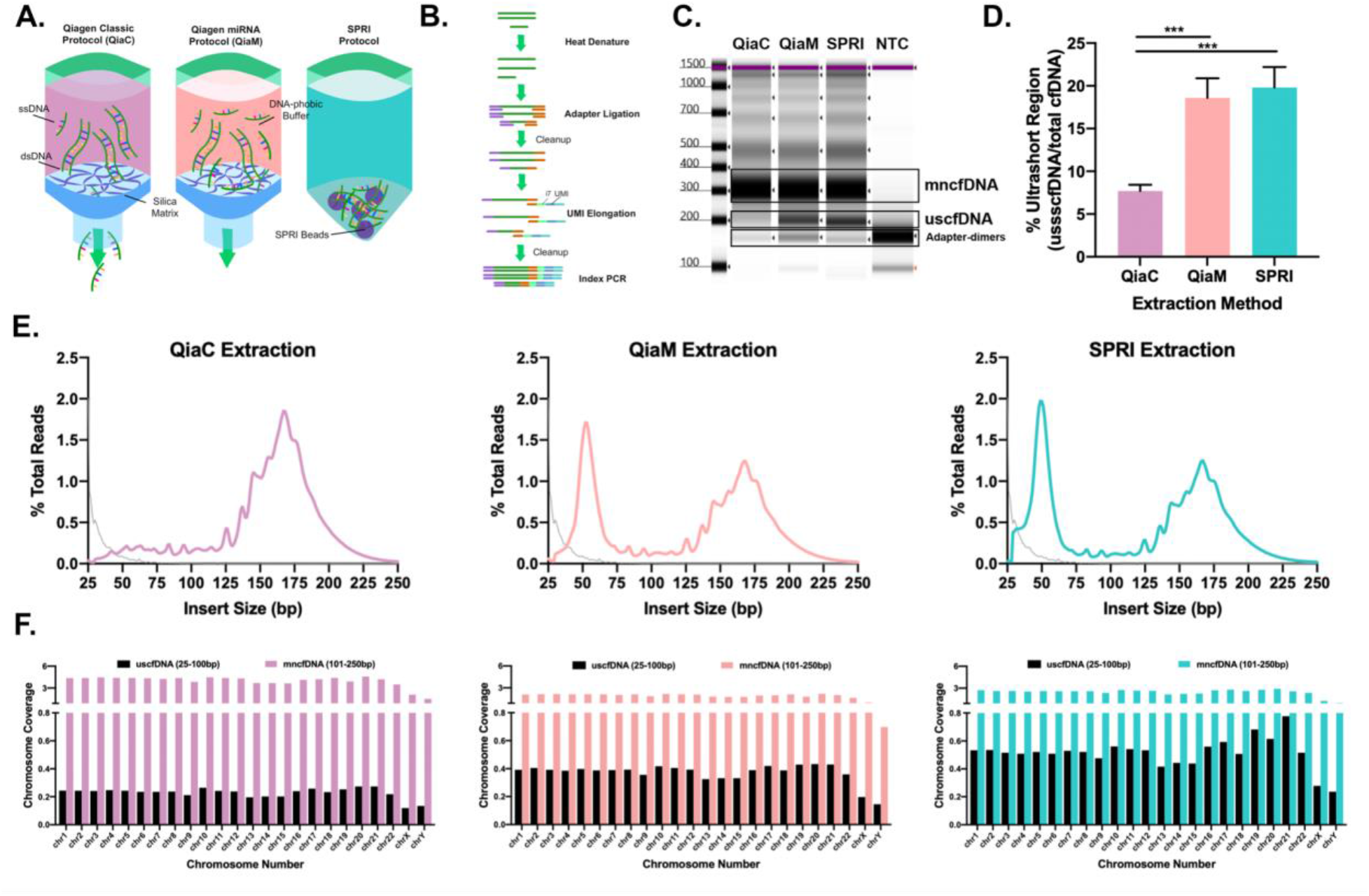
uscfDNA-seq reveals a population of ultrashort cfDNA fragments at 50nt in plasma of healthy donors. **A**. Schematic of the three extraction protocols compared in this manuscript. QiaM and SPRI protocols utilize an enhanced DNA-phobic buffer in order to retain the low-molecular nucleic acids for downstream analysis (refer to methods for details). **B**. Single-stranded library preparation can incorporate dsDNA, ssDNA, and nicked DNA into the library. Unique molecular identifiers (UMI) are incorporated during the library preparation to remove PCR duplicates. **C**. uscfDNA-seq using QiaM or SPRI reveal a distinct uscfDNA band at 200bp (∼50bp after adapter dimer subtraction) compared to QiaC. **D**. QiaM and SPRI extraction method can reproducibly isolate the 200bp fragment in ten healthy donors based on quantification of electrophoresis output (200bp band divided by (200bp + 300bp). **E**. Alignment of sequenced reads from QiaM and SPRI extracted samples exclusively show the uscfDNA at 50bp in addition to the mncfDNA peak at ∼160bp. Extraction methods: QiaC (fuschia), QiaM (pink), and SPRI (teal). Grey line represents sequencing of no template control. **F**. The uscfDNA population of the QiaM and SPRI map along the genome. *** *p* < 0.001. The paired two-tailed student-test test was performed after ANOVA analysis. Bars graphs represent standard error of Mean (SEM).

Similarly, we observed that using the miRNA protocol of a commercial silica column-based extraction kit (QiaM) which incorporates higher isopropanol volume will also enhance the capture of low-molecular nucleic acids **(Supp. Figure 1A**). Interestingly, the miRNA purification protocol is associated with slower flow through the silica column. SEM images of the silica column indicate a reduction in pore size accompanied by sheet-like deposits possibly derived from increased isopropanol precipitation of organic matter in the plasma (**Supp. Fig 1B**).

Single-stranded libraries **(Figure 1B)** were made from cell-free DNA extracted by QiaM and SPRI methods which revealed a distinct cfDNA band at 200bp in the electropherogram corresponding to about 50bp of insert size (the library preparation adds about 150 bp-worth of adapters) compared to the standard protocol of the commercial silica column-based extraction kit (QIAGEN QIAamp Circulating Nucleic Acid Kit, or QiaC) **(Figure 1C and 1E)**. In all three extraction methods the mncfDNA peak (300bp before adapter removal) is present. Treatment with RNase Cocktail digestion prior to library preparation did not appreciably decrease the uscfDNA band ruling out the presence of RNA **(Supp. Figure 1C)**. This is a reproducible phenomenon with similar observations in multiple donors **(Figure 1D and Supp. Figure 2A)**, and irrespective of the whole blood collection tube used (**Supp. Figure 1D**, and see below). Extractions performed on TE buffer alone did not manifest any uscfDNA or mncfDNA bands except for adapter-dimer bands **(Supp. Figure 2B)**, ruling out the presence of contaminants.

Upon sequencing and alignment to the human genome, the cell-free DNA fragment sizes were divided into two distinct size populations (25-100bp and 101-250bp) with QiaM and SPRI both showing increased coverage of the ultrashort population **(Figure 1E)**. There was approximately even coverage across the genome of the ultrashort population **(Figure 1F)**. It has been previously reported that mitochondria-derived cell-free DNA is fairly short (50bp) but we found that it only contributed a minority (<0.1%) of the total mappable DNA reads **(Supp. Figure 2C)** despite the fact that QiaM and SPRI methods enrich for ultrashort fragments (**Supp. Figure 2D)**.

Plasma from K2EDTA vacu-containers contained uscfDNA **(Figure 1)** but it has been reported to be associated with cell-free DNA degradation.^8^ Thus, to rule out the possibility of uscfDNA as an artifact of sample collection, StreckDNA tubes (the gold-standard for cell-free DNA preservation due to their ability to decrease white blood cell rupture and subsequent genomic DNA contamination in the sample) was also tested for presence of uscfDNA. An alternative, StreckRNA, which is used to preserve RNA (a low molecular nucleic acid) and exosomes was also tested. All three collection tubes allowed us to detect the presence of the uscfDNA population (**Supp. Fig 1D**).

To examine the properties of strandedness, treatment of the extracted DNA by a dsDNA specific enzyme (dsDNase) prior to library preparation showed a marked reduction in the mncfDNA band (300bp) while preserving the uscfDNA band (200bp) (**Figure 2A and Supp. Fig 3A)** in both QiaM and SPRi isolated samples. In contrast, digestion by single-strand specific nucleases (S1, exo 1, and P1) showed significant reduction in the uscfDNA band while preserving the mncfDNA band in both plasma extracted by the QiaM and SPRI protocol. Sequencing and alignment of these libraries confirmed the results from the electropherograms **(Figure 2A, bottom panels)**. These results strongly indicate the single-stranded nature of the uscfDNA.

**Figure. 2.**
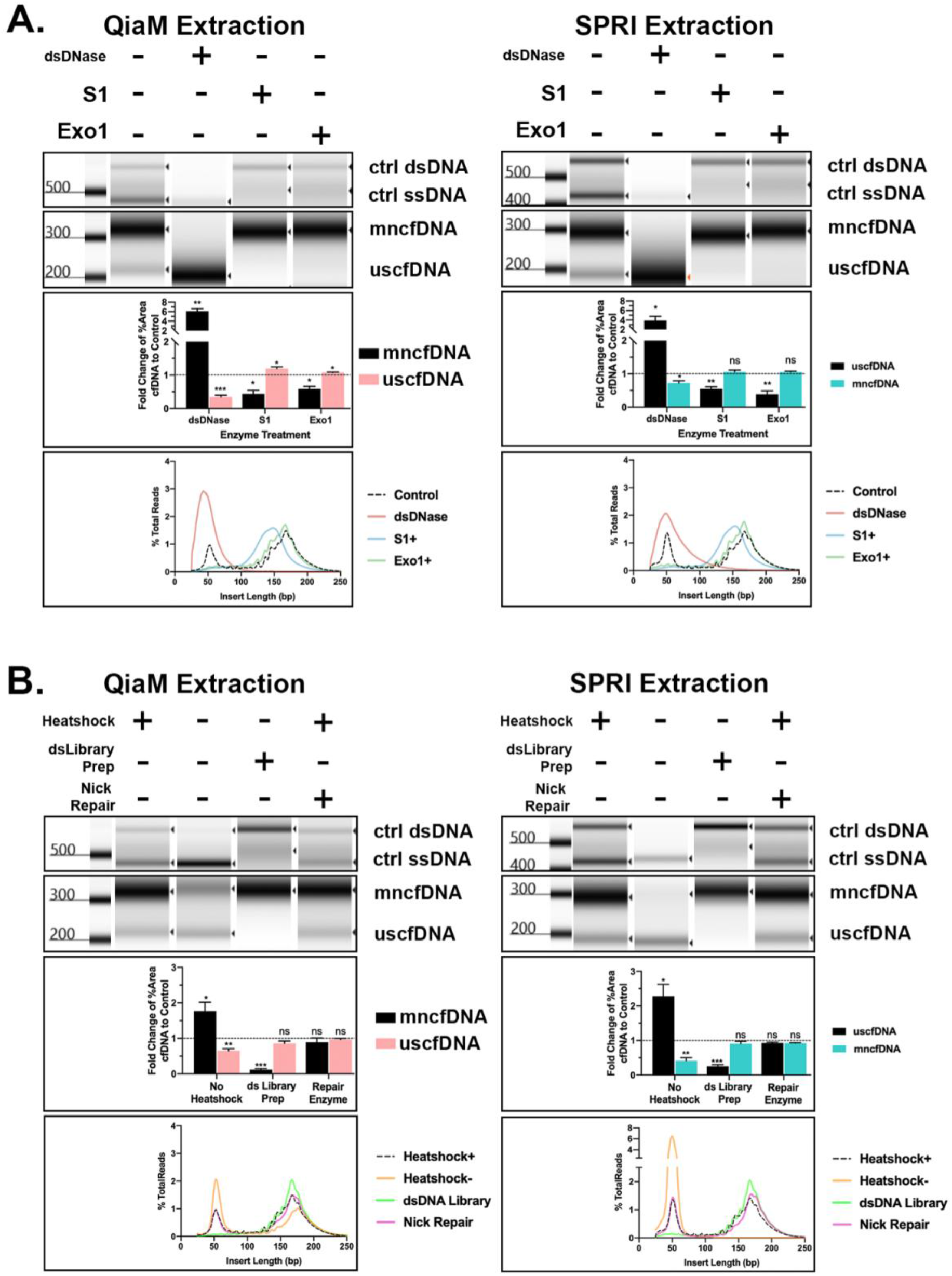
uscfDNA population is predominantly single-stranded. **A**. Compared to control (Heatshock+) digestion of extracted cfDNA by both QiaM (Pink) and SPRI (Teal) prior to library preparation using dsDNA-specific dsDNase only reduces the mncfDNA population but retains the uscfDNA bands. In contrast, ssDNA-specific nucleases (S1 and Exo1) show significant reduction of the uscfDNA band. **B**. Absence of heat-denaturation (heatshock) at the beginning of the single-stranded library preparation reduces mncfDNA incorporation but not the uscfDNA population. Opposingly, dsDNA library preparation only manifests the mncfDNA band. Pre-treatment with the DNA nick-repairing enzyme does not reduce the uscfDNA population. Alignment of sequenced libraries reflect observations in electropherograms. Reads with insert size under 25bp and above 250 were excluded from the plots. Bar graphs composed of three different healthy donors. The paired two-tailed student-test test was performed after ANOVA analysis. * *p* < 0.05, ** *p* < 0.01, and *** *p* < 0.001. Lambda-sequenced 460bp dsDNA and 356nt ssDNA were used as positive controls. Adapter-dimers have been cropped from the presented electropherograms. Bars graphs represent standard error of Mean (SEM).

To corroborate the single-stranded nature of this DNA we leveraged the differences in the adapter ligation chemistry between ssDNA and dsDNA library kits **(Figure 2B)**. The uscfDNA peak was absent in the dsDNA library preparation (which only incorporates intact double-stranded structures) suggesting that the ultrashort population is endogenously single-stranded in nature. By contrast, the ssDNA library kits require initial heat denaturation (98°C for 3 minutes) to efficiently incorporate adapter molecules into the library. By skipping this step, the presence of the 200bp population remained suggesting that the uscfDNA population is single-stranded **(Figure 2B)**. Finally, to determine if the source of the uscfDNA derived from nicked dsDNA, we pretreated the extracted nucleic acids with a nick repair enzyme but did not observe a reduction of ultrashort fragments in the final library. This suggests that the uscfDNA are not derived from nicked mncfDNA. These observations were consistent among three replicates **(Supp. Fig 3AB)**.

Alignment of sequenced digestion libraries recapitulated the findings previously mentioned with some interesting observations (**Figure 2A, B and Suppl. 4AB**). Firstly, the S1 treated samples showed a 10bp downshift in the modality of the mncfDNA peak (from 160 to 150bp). Secondly, both the S1 and nick-repair enzyme treatment flattened the periodicity on the left side of the mncfDNA peak. These observations suggest that the 10bp periodicity may be a result of nicked mncfDNA at certain fragment lengths.^2^ The S1 enzyme may also be digesting jagged edges flanking the mncfDNA. These observations are worth exploring in depth in future studies.

In order to compare the efficiency of the extraction methods, non-human ssDNA oligos designed from the *E. coli* phage lambda genome of sizes 30, 50, 75, 100, 150, and 200nt **(Suppl. Table 2)** were spiked into the plasma prior to extraction and library preparation. The uscfDNA extraction methods (QiaM and SPRI) retain ultrashort fragments in plasma with greater efficiency compared to the regular QiaC protocol **(Suppl. Fig 5A and B)**. Interestingly, the SPRI extraction method showed improved retention of 30 and 50nt ssDNA compared to QiaM. Although these two extraction methods show improved ability in retaining low-molecular ssDNA, their yield suggests that there is still substantial loss. Hence, further refining of future methods to improve the yield is warranted. Advantages of the current bead-based methods is that they limit physical loss of ultrashort cfDNA fragments compared to silica columns that utilize flow through the pores. However, the observed presence of adapter-dimers is suggestive of the presence of inhibiting factors that may interfere with downstream enzyme activity.

The functions of RNA, a prominent single-stranded entity, are well described. RNA is involved in transcription, amino-acid transfer, protein-complexes, gene expression, and signal-transfer via exosomes. By comparison, circulating ssDNA biology has been largely unexplored, and it is plausible that ssDNA may have more functions than initially thought. In molecular biology, there is limited technology to evaluate ssDNA. In this report we demonstrate the **uscfDNA-seq** pipeline which can use two cell-free DNA extraction methods, one modified commercial and one customized, with a single-stranded library preparation to reveal the presence of a unique class of ultrashort single-stranded cell-free DNA of nuclear origin with a modal size of 50nt. The use of **uscfDNA-seq** suggests that with regards to cfDNA liquid biopsy, the uscfDNA population must be considered, in conjunction with conventional mncfDNA, for biomarker identification and diagnosis. We have previously found evidence that ultrashort circulating tumor DNA contained in plasma from non-small cell lung carcinoma patients can also harbor mutations corresponding to the mncfDNA and tissue genotyping.^9^ However, the pipeline was not optimized for single-strand DNA, in contrast to the methodology we present here. Therefore, careful examination of uscfDNA may likely provide new opportunities in molecular diagnostics and cfDNA biology in the future.

## Methods

### Clinical Samples

Plasma from healthy donors was commercially purchased from Innovative Research (IPLASK2E10ML). One donor provided whole blood collected into three vacutainers, K2EDTA, StreckDNA, and StreckRNA (Streck, 218961 and 230460). According to vendor instructions, whole blood was spun at 5000xG for 15 minutes and plasma was removed using a plasma extractor. Age and gender of the donors can be found in the supplemental chart **(Suppl. Chart 1)**.

### Nucleic Acid Extraction

1 mL of plasma was extracted with three different methods. Using the QIAmp Circulating Nucleic Acid Kit (Qiagen, 55114) we followed two of the manufacturer protocol: Purification of Circulating Nucleic Acids from 1mL of Plasma (QiaC) and Purification of Circulating microRNA from 1ml of Plasma (QiaM). Proteinase-K digestion was carried out as instructed. Carrier RNA was not used. The ATL Lysis buffer (Qiagen, 19076) was used as indicated in the microRNA protocol. The final elution volume was 40µl.

In the magnetic bead-based uscfDNA extraction, 100µL of Proteinase K (20mg/mL, Zymogen, D3001-2-1215) and 56µL 20% SDS (Invitrogen, AM9820) was added to 1mL of human plasma and incubated for 30minutes at 60°C. After cooling to ambient room temperature, 540µL SPRI-select beads (Beckman Coulter, B22318) and 3000µL of 100% isopropanol (Fisher, BP26181) were added to the plasma and incubated for 10 minutes on the benchtop. The plasma was then centrifuged at 4000xG for five minutes. The supernatant was removed and discarded. The pellet was resuspended using 1mL of 1x TE Buffer (Invitrogen, AM9848) and divided into 500µl aliquots into two phase lock tubes (Quantabio, 10847-802). An equal volume (500µL) of phenol:chloroform:isoamyl alcohol with equilibrium buffer was added (Sigma, P2069-100mL) and contents were vortexed for 15 seconds. The tubes were then centrifuged at 19000xG for five minutes. This was repeated twice (vortexed and centrifuged). The upper clear supernatant was pipetted and transferred to a 15mL conical tube SPRI-select beads and 3000µL of 100% isopropanol were added to the plasma and incubated for 10 minutes on the benchtop. The tube was placed on a magnetic rack for five minutes to allow for the beads to migrate. The supernatant was discarded and the beads were washed twice with 5ml of 85% ethanol. Once the second ethanol wash was removed the beads were left to air dry for 10minutes. The beads were then resuspended in 30µL of elution buffer (Qiagen, 19086) and incubated for 2 minutes. After the beads were transferred to a 1.5mL tube and magnet rack to separate the beads. Once the solution was clear (∼2 minutes) the 30µL of elution was transferred to another 1.5mL tube and combined with 1µL of 20mg/ml glycogen (Thermo, R0561), 44µL of 1xTE Buffer, 25µL of 3M sodium acetate (Quality Biological INC, 50-751-7660), 250µL of 100% ethanol and placed at -80°C overnight. The tube was then centrifuged at 19000xG for 15 minutes. The supernatant was removed and replaced with 200µL of 80% ethanol. This was done 2 more times. The supernatant was removed and the pellet was resuspended in a 30µL of elution buffer and combined with 90µL of SPRI-select beads, 90µL of 100% isopropanol and incubated for 10 minutes. The tube was placed on a magnetic rack for five minutes to allow for the beads to migrate. The supernatant was discarded and the beads were washed twice with 200µL of 80% ethanol. Once the second ethanol wash was removed the beads were left to air dry for 10minutes. The beads were then resuspended in 40µL of Qiagen elution buffer.

### Library Preparations

Single-stranded DNA library preparation was performed using the SRSLY™ PicoPlus DNA NGS Library Preparation Base Kit with the SRSLY 12 UMI-UDI Primer Set, UMI Add-on Reagents, and purified with Clarefy Purification Beads (Claret Bioscience, CBS-K250B-24, CBS-UM-24, CBS-UR-24, CBS-BD-24). Since there is currently no optimized method to measure ussscfDNA, 18µL of extracted cfDNA was used as input and heat-shocked as instructed. To retain a high proportion of small fragments the low molecular weight retention protocol was followed for all bead-clean up steps. The index reaction PCR was run for 11 cycles. For double-stranded DNA libraries the NEB Ultra II (New England Bio, E7645S) was used with an 9µL aliquot of extracted cfDNA according to the manufacturer’s instructions with some modifications: the adapter ligation was performed using 2.5 µl of NEBNext® Multiplex Oligos for Illumina (Unique Dual Index UMI Adaptors RNA Set 1 - NEB, cat# E7416S); the post-adapter ligation purification was performed using 50 µl of purification beads and 50 µl of purification beads’ buffer, while the second (or post-PCR) purification was performed using 60 µl of purification beads (to retain smaller fragments). The PCR was performed using the MyTaq HS mix (Bioline, BIO-25045) - or the Q5 polymerase (NEB, M0491S) for 10 PCR cycles.

### Sequencing

Final library concentrations were measured using the Qubit Fluorometer (Thermo, Q33327) and quality assessed using the Tapestation 4200 using D1000 High-Sensitivity Tapes (Agilent, G2991BA and 5067-5584). Final libraries were run on Nova-Seq SP 300 (150×2).

### Bioinformatic Processing

Sequence reads were demultiplexed using SRSLYumi (SRSLYumi 0.4 version, Claret Bioscience), python package. Fastq files were trimmed with (fastp,^10^ using adapter sequence AGATCGGAAGAGCACACGTCTGAACTCCAGTCA (r1) and AGATCGGAAGAGCGTCGTGTAGGGAAAGAGTGT (r2) and a Phred score of >15. Then sequenced reads were aligned against the human reference genome [GenBank:GCA_000001305.2] and LambdaPhage Genome [GeneBank:GCA_000840245.1] using BWA-mem.^11^ The duplicated reads were removed using Picard Toolkit (http://broadinstitute.github.io/picard/), after sorting and filtering with samtools (samtools 1.9 version).^12^ Quality control was performed with Qualimap Version 2.2.2c.^13^

### Nuclease Digestions for Analysis of Strandedness

Prior to library preparation, the extracted cfDNA was digested with various strand-specific nucleases. For all reactions **500pg** of control oligos (350nt ssDNA and 460bp dsDNA lambda sequence, IDT) was spiked into 20µL of cfDNA. DNA was purified by combining 30µL of reaction buffer and combined with 90µL of SPRI-select beads, 90µL of 100% isopropanol and incubated for 10 minutes. The tube was placed on a magnetic rack for five minutes to allow for the beads to migrate. The supernatant was discarded and the beads were washed twice with 200µL of 80% ethanol. Once the second ethanol wash was removed the beads were left to air dry for 10minutes. The beads were then resuspended in 20µL of Qiagen elution buffer.

Non-strand specific DNA digestion: 20µL cfDNA was combined with 1µL **DNase I** (Invitrogen, 18-068-015), 3µL 10xDNase 1 Buffer, 6µL of ddH2O incubated for 15minutes at 37°C and heat inactivated for 15 minutes at 80°C with 1µL of 0.5M EDTA.

ssDNA-specific Digestion: 20µL cfDNA was combined with 1µL 1x **S1** (Thermo, EN0321), 6µL 5x S1 Buffer, 3µL of ddH2O incubated for 30 minutes at room temperature and heat inactivated for 15 minutes at 80°C with 2µL of 0.5M EDTA.

ssDNA-specific Digestion: 20µL cfDNA was combined with 1µL 0.1x **P1** (NEB, M0660S), 3µL NEBuffer r1.1, 6µL of ddH2O incubated for 30 minutes at 37°C and inactivated with 2µL of 0.5M EDTA.

ssDNA-specific Digestion: 20µL cfDNA was combined with 3µL **Exonuclease 1** (NEB, M0293S), 3µL 10x Exo 1 Buffer, 4µL of ddH2O incubated for 30 minutes at 37°C and heat inactivated for 15 minutes at 80°C with 1µL of 0.5M EDTA.

dsDNA-specific Digestion: 20µL cfDNA was combined with 2µL **dsDNase** (ArcticZyme, 70600-201), 8µL of ddH2O incubated for 30 minutes at 37°C and heat inactivated for 15 minutes at 65°C with 1mM DTT.

Nick Repair Analysis: 20µL cfDNA was combined with 1µL **PrePCR Repair** (NEB, M0309S), 5µL ThermoPol Buffer (10x), 0.5µL of NAD+ (100x), 2µL of Takara 2.5mM dNTP, 21.5 ddH2O incubated for 30 minutes at 37°C and placed on ice.

RNA Digestion: 20µL of cfDNA was combined with 1µL of **RNase Cocktail** (Thermo, AM228). For 20 minutes at 30°C prior to input into the library preparation.

### ssDNA Ladder to Determine Efficiency

2ng ssDNA ladder of various sizes (30-200) was spiked in 1mL healthy plasma prior to extraction. Final elution was 40µL and 18µL was used for each final library. Oligonucleotides were manufactured by a commercial vendor (IDT, Custom Order).

### Scanning electron microscope (SEM)

After processing PBS or plasma samples with QiaC or QiaM protocol, the columns were air-dried at room temperature. They were cut into proper height to expose the membrane and fitted to the sample stage. The samples were coated with platinum and the detailed morphology of the membrane was examined by Focus-Ion Beam/Scanning Electron Microscopy(FEI, Nova 200 NanoLab).

## Acknowledgements

This work was supported by NIH grants UH2/UH3 CA206126, UH2/UH3 TR000923, UO1 CA233370, Spectrum Solutions 20212918 (DTWW), and the Canadian Institute of Health Research Doctoral Foreign Study Award, Tobacco Related Disease Research Program (TRDRP) Predoctoral Fellowship, Jonsson Comprehensive Cancer Center Predoctoral Fellowship, and the NIH grant UL1TR001881 (JC).

## Author Contributions

Conceptualization, J.C., M.M., M.P., and D.T.W.W.; Data Curation, J.C., M.M., W.H., and T.P.; formal analysis, J.C., M.M., Y.H.; funding acquisition, J.C. and D.T.W.W.; investigation, J.C., M.M., M.P., and D.T.W.W.; methodology, J.C., M.M., M.P., and D.T.W.W.; project administration, M.P. and D.T.W.W.; Resources, D.T.W.W; software, J.C., M.M., ; supervision, M.P. and D.T.W.W; visualization, J.C., Y.H., and M.M.; writing— original draft, J.C., M.M..; writing—review and editing, J.C., M.M., W.H., Y.H., T.P., F.L., F.W., D.C., H.J., K.C., W.S., M.P., and D.T.W.W. All authors have read and agreed to the submitted version of the manuscript.

