## Supplemental Figures and Tables for "Plasma Contains Ultra-short Single-stranded DNA in Addition to Nucleosomal cfDNA"

### Supplementary Figures and Tables

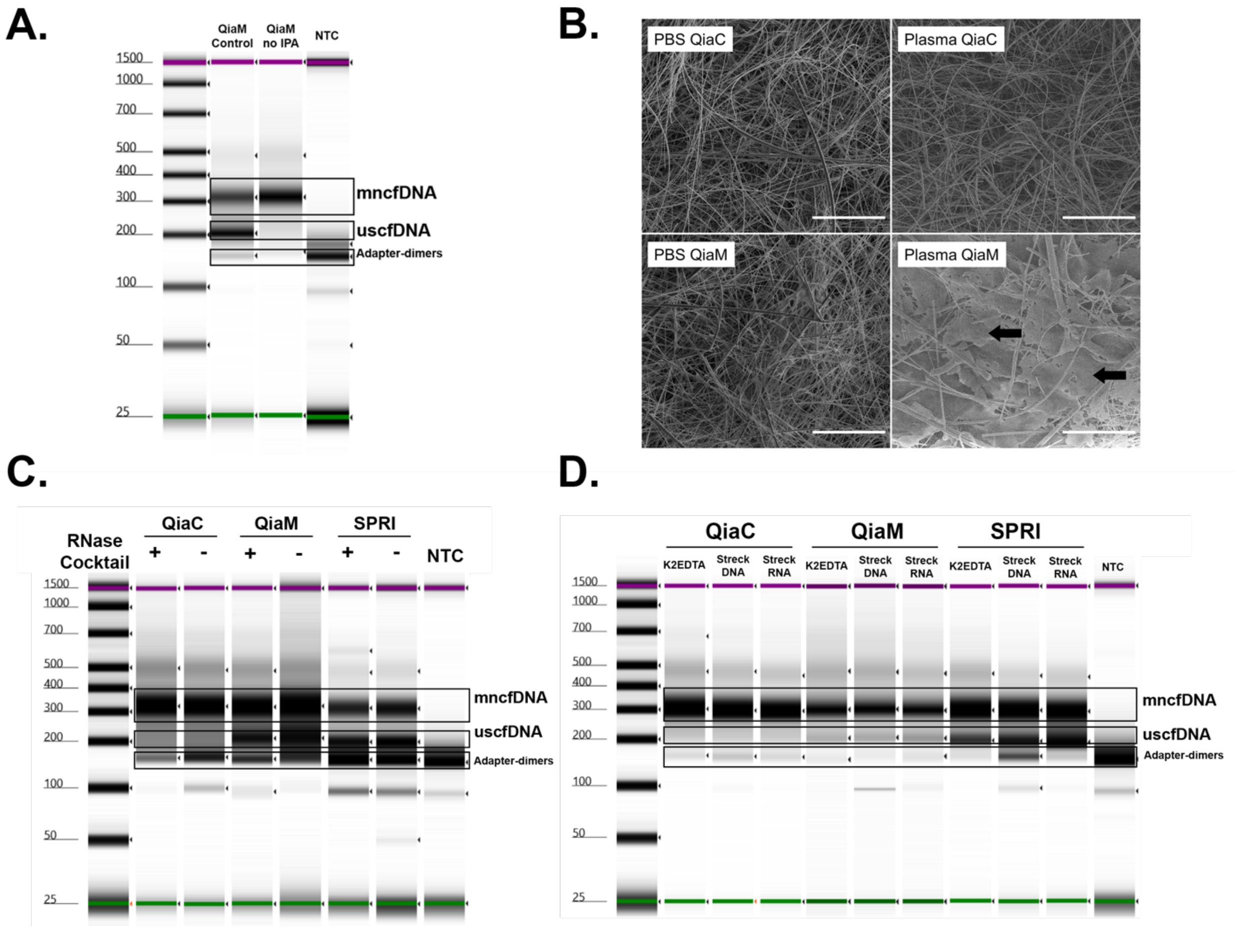


**Supplemental Figure 1 | A**. RNase cocktail digestion prior to library preparation does not reduce the uscfDNA band in QiaM and SPRI extracted samples. **B.** SEM images of the Qiagen silica filter show global sheet-like deposits (black arrows) only in QiaM extraction of plasma. Scale bars (white line) represents 50um. **C.** The increased isopropanol (1.8ml to 2.3ml) is integral to retaining the uscfDNA from plasma. **D.** UscfDNA exists independently of the whole blood collection tube.

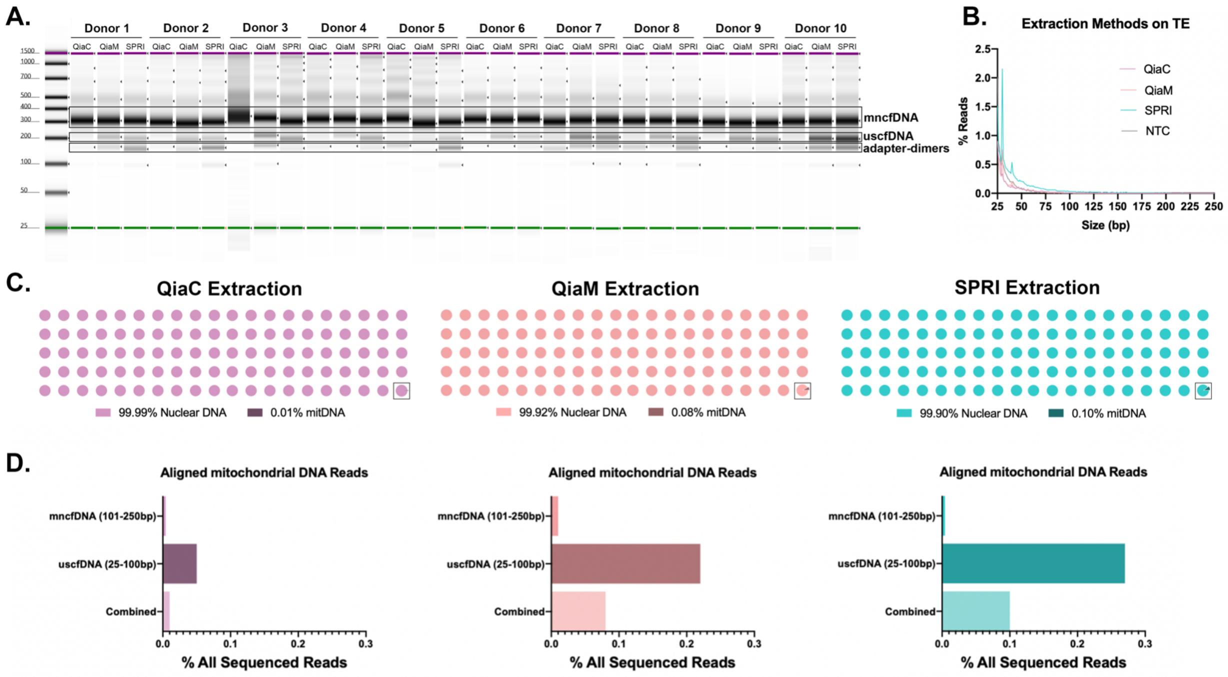


**Supplemental Figure 2 | A.** Electropherogram images of ten healthy donors extracted with QiaC, QiaM, and SPRI showing the presence of uscfDNA **B**. TE buffer control extracted with the three methods do not produce uscfDNA or mncfDNA peaks when aligned to the human genome. **C.** The majority of DNA aligns to the nuclear genome and not to the mitochondrial genome. Square indicates the visual representation of contribution of mitochondria reads. Extraction methods: QiaC (fuschia), QiaM (pink), and SPRI (teal). **D.** QiaM and SPRI are enriched for mitochondrial DNA in the uscfDNA population but still is a minor fraction of total DNA.

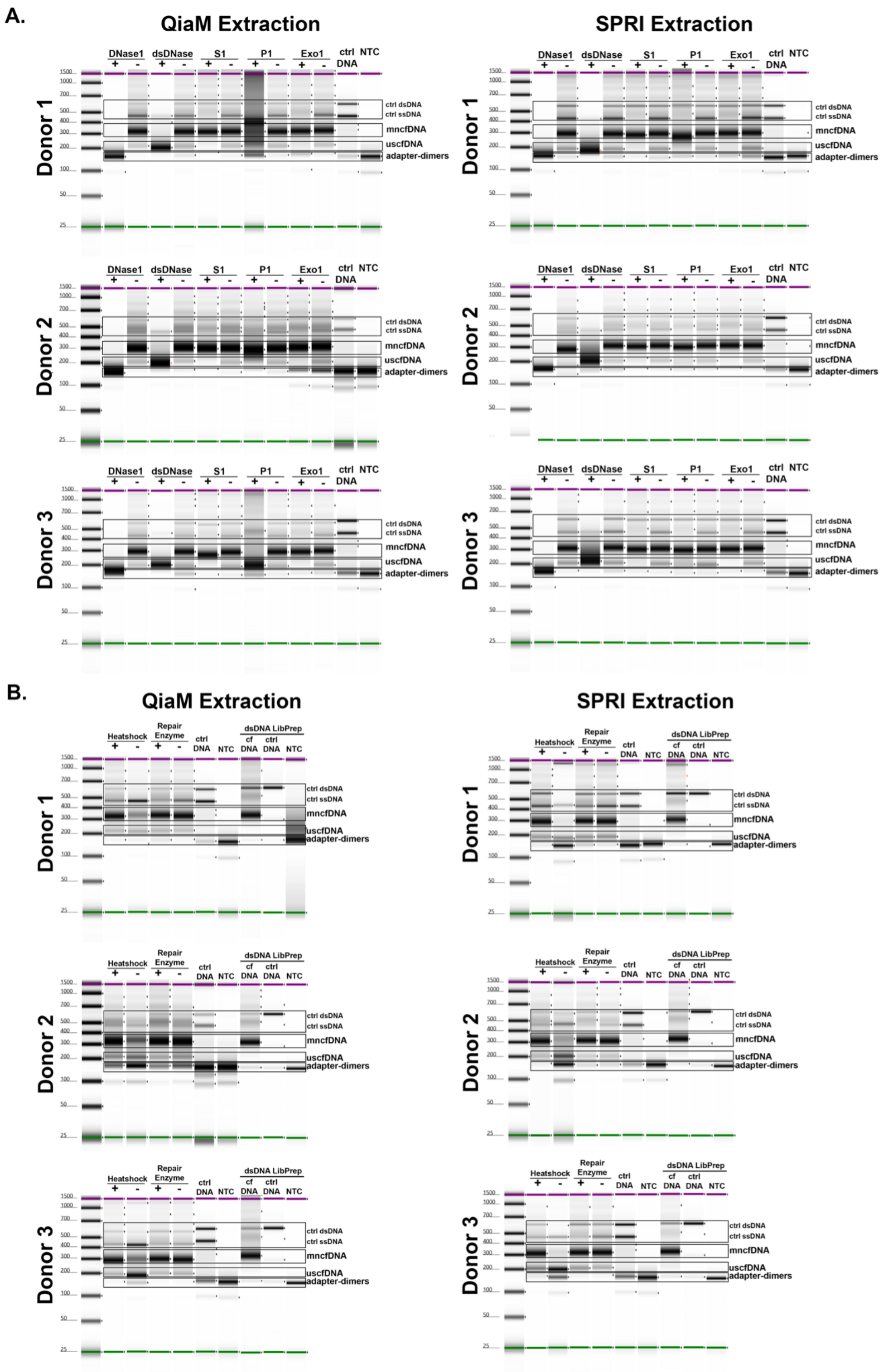
A.
QiaM Extraction
SPRI Extraction
DNase1
dsDNase
S1
P1
Exo1
ctrl DNA
NTC
+
-
Donor 1
Donor 2
Donor 3
ctrl dsDNA
ctrl ssDNA
mncfDNA
uscfDNA
adapter-dimers
1500
1000
700
500
400
300
200
100
50
25
B.
QiaM Extraction
SPRI Extraction
Heatshock
Repair Enzyme
ctrl DNA
NTC
dsDNA LibPrep
cf DNA
ctrl DNA
NTC
ctrl dsDNA
ctrl ssDNA
mncfDNA
uscfDNA
adapter-dimers

**Supplemental Figure. 3 | A.** Electropherograms from nuclease digestions of extracted cfDNA prior to library preparation. **B.** Electropherograms from cfDNA undergoing ssDNA, dsDNase library preparation and nick-repair enzyme treatment. Replicate experiments using plasma from three healthy donors extracted by QiaM and SPRI.

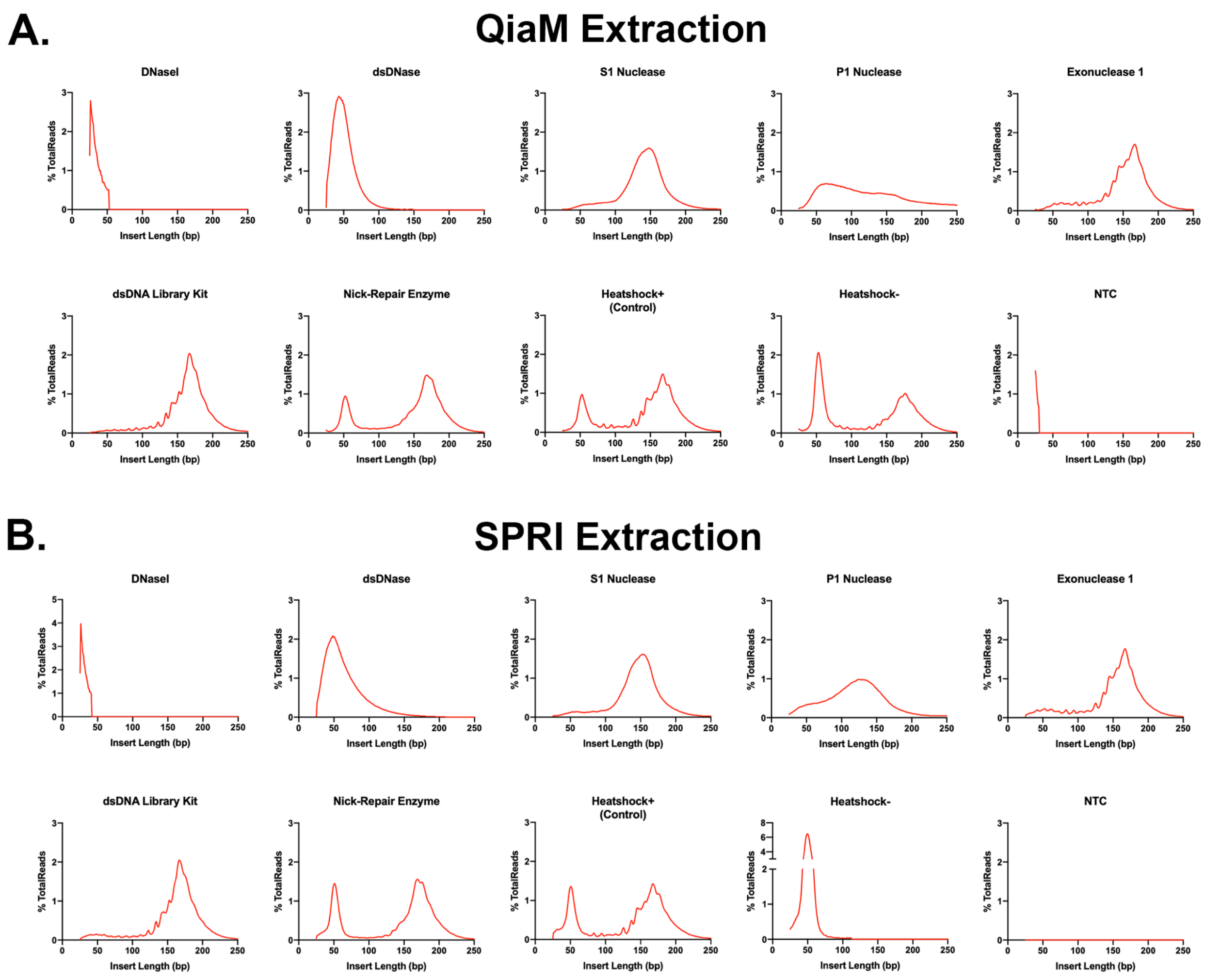


**Supplemental Figure. 4 |** Alignment of sequenced libraries to human genome pretreated by digestions and library preparation variations from Donor 1 of Sup Fig 3 extracted by QiaM (**A**) and SPRI (**B**). Reads with insert size under 25bp and above 250bp were excluded from the plots.

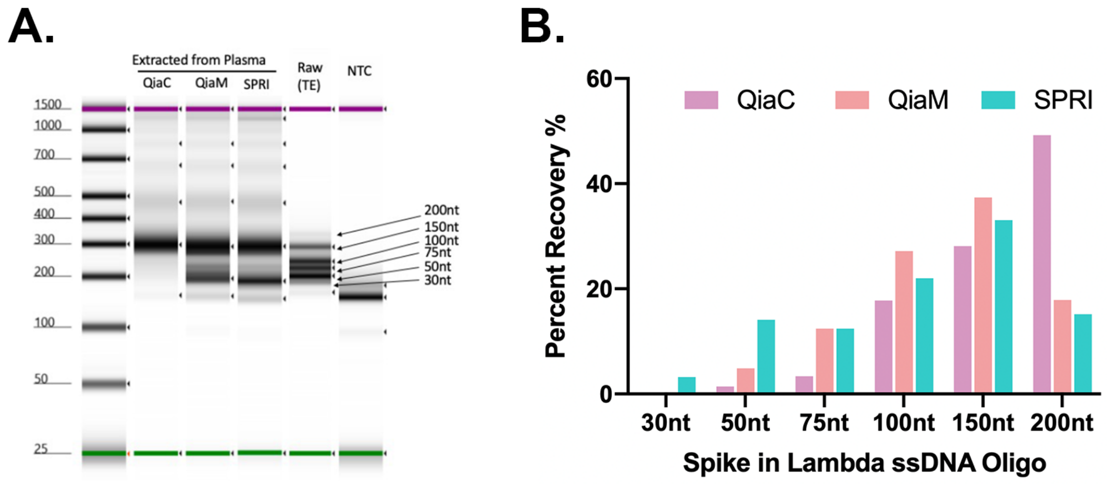


**Supplemental Figure. 5 | A.** Extraction of healthy plasma spiked with a ladder of lambda ssDNA oligos show various retention efficiencies between QiaC, QiaM and SPRI methods. **B.** After alignment to the lambda genome shows QiaM and SPRI methods have greater efficiency of extracting ultrashort ssDNA.

**Supplemental Table 1 | Healthy Plasma Donor Information**

| Assay | Gender | Age |
|---|---|---|
| Digestions Donor 1 | Male | 47 |
| Digestions Donor 2 | Female | 57 |
| Digestions Donor 3 | Male | 35 |
| Healthy 10 Replicate Donor 1 | Male | 45 |
| Healthy 10 Replicate Donor 2 | Male | 18 |
| Healthy 10 Replicate Donor 3 | Male | 23 |
| Healthy 10 Replicate Donor 4 | Male | 26 |
| Healthy 10 Replicate Donor 5 | Male | 38 |
| Healthy 10 Replicate Donor 6 | Male | 33 |
| Healthy 10 Replicate Donor 7 | Male | 22 |
| Healthy 10 Replicate Donor 8 | Male | 37 |
| Healthy 10 Replicate Donor 9 | Male | 27 |
| Healthy 10 Replicate Donor 10 | Male | 41 |

### Supplemental Table 2 | Synthetic Oligomers and Primers

<table>
<tr><th>Name</th><th>Size</th><th>ss/ds</th><th>Lambda phage region</th><th>Notes</th></tr>
<tr><td>Lambda dsDNA Control</td><td>459 bp</td><td>ds</td><td>27’944:28’402</td><td>PCR product, no UMI</td></tr>
<tr><td colspan="5">5′-CAAACTGCGCAACTCGTGAAAGGTAGGCGGATCCCCTTCGAAGGAAAGACCTGATGCTTTTCGTGCGCGCATAAAATACCTTGATACTGTGCCGGATGAAAGCGGTTCGCGACGAGTAGATGCAATTATGGTTTCTCCGCCAAGAATCTCTTTGCATTTATCAAGTGTTTCCTTCATTGATATTCCGAGAGCATCAATATGCAATGCTGTTGGGATGGCAATTTTTACGCCTGTTTTGCTTTGCTCGACATAAAGATATCCATCTACGATATCAGACCACTTCATTTCGCATAAATCACCAACTCGTTGCCCGGTAACAACAGCCAGTTCCATTGCAAGTCTGAGCCAACATGGTGATGATTCTGCTGCTTGATAAATTTTCAGGTATTCGTCAGCCGTAAGTCTTGATCTCCTTACCTCTGATTTTGCTGCGCGAGTGGCAGCGACATGGTTTGTTGT-3′</td></tr>
<tr><td>Lambda ssDNA Control</td><td>350 nt</td><td>ss</td><td>7’582:7’930</td><td>IDT synthesized</td></tr>
<tr><td colspan="5">5′-CCTGGCCAGAATGCAATAACGGGAGGCGCTGTGGCTGATTTCGATAACCTGTTCGATGCTGCCATTGCCCGCGCCGATGAAACGATACGCGGGTACATGGGAACGTCAGCCACCATTACATCCGGTGAGCAGTCAGGTGCGGTGATACGTGGTGTTTTTGATGACCCTGAAAATATCAGCTATGCCGGACAGGGCGTGCGCGTTGAAGGCTCCAGCCCGTCCCTGTTTGTCCGGACTGATGAGGTGCGGCAGCTGCGGCGTGGAGACACGCTGACCATCGGTGAGGAAAATTTCTGGGTAGATCGGGTTTCGCCGGATGATGGCGGAAGTTGTCATCTCTGGCTTGGAC-3′</td></tr>
<tr><td>lambda 200</td><td>198 nt</td><td>ss</td><td>12’051:12’248</td><td>IDT synthesized, internal-UMI 12nt</td></tr>
<tr><td colspan="5">5′-AAGGCGGAGAGTCAGTTCGCGGNNNNNNNNNNNNCGGCGCAACGTCGCCAGCTGTCTGCACAGGAGAAATCCCTGCTGGCGCATAAAGATGAGACGCTGGAGTACAAACGCCAGCTGGCTGCACTTGGCGACAAGGTTACGTATCAGGAGCGCCTGAACGCGCTGGCGCAGCAGGCGGATAAATTCGCACAGCAGCAA-3′</td></tr>
<tr><td>lambda 150</td><td>150 nt</td><td>ss</td><td>35’073:35’201+UMI</td><td>IDT synthesized, 3’-UMI 12nt</td></tr>
<tr><td colspan="5">5′-GCGTCCACTGCATGTTATGCCGCGTTCGCCAGGCTTGCTGTACCATGTGCGCTGATTCTTGCGCTCAATACGTTGCAGGTTGCTTTCAATCTGTTTGTGGTATTCAGCCAGCACTGTAAGGTCTATCGGATTTAGTGCNNNNNNNNNNNN-3′</td></tr>
</table>

| | | | | |
|---|---|---|---|---|
| lambda 100 | 100 nt | ss | 41’091:41’178+UMI | IDT synthesized, 3’-UMI 12nt |
| 5’-<br>TCGTTAGTTTCTCCGGTGGCAGGACGTCAGCATATTTGCTCTGGCTAATGGAGCAAAAGCGACGGGCAGGTAA<br>AGACGTGCATTACGTNNNNNNNNNNNN-3’ | | | | |
| lambda 75 | 75 nt | ss | 18’204:18’266+UMI | IDT synthesized, 3’-UMI 12nt |
| 5’-<br>TCGTATCGCATTTATTGACCCGGCAAACGGGAATGAAACGCCGATGTTTGTGGCGCAGGGCAANNNNNNNNNN<br>NN-3’ | | | | |
| lambda 50 | 50 nt | ss | 2’321:2’359+UMI | IDT synthesized, 3’-UMI 12nt |
| 5’-ACCGCTTCCCGGTGCCGTTCACTTCCCGAATAACCCGGANNNNNNNNNNNN-3’ | | | | |
| lambda 30 | 30 nt | ss | 4’278:4’300+UMI | IDT synthesized, 3’-UMI 9nt |
| 5’-ACGCGGTGACGACTATCAGGAAANNNNNNN-3’ | | | | |
| I7 Extension Primer Sequence (i7 ext) | 75 nt | 5’-<br>CAAGCAGAAGACGGCATACGAGATNNNNNNNNNXXXXXXXXGTGACTGGAG<br>TTCAGACGTGTGCTCTTCCGATCT-3’ | | |
| Forward Index Primer Sequence (i5) | 70 nt | 5’-<br>AATGATACGGCGACCACCGAGATCTACACXXXXXXXXACACTCTTTCCCTA<br>CACGACGCTCTTCCGATCT-3’ | | |
| Reverse Index Primer SEquence (Ui7) | 21 nt | 5’- CAAGCAGAAGACGGCATACGA-3’ | | |

### Supplemental Table 3 | Numerical values for Figure 1D.

| uscfDNA region/total cfDNA (uscfDNA + mncfDNA) | Donor 1 | Donor 2 | Donor 3 | Donor 4 | Donor 5 | Donor 6 | Donor 7 | Donor 8 | Donor 9 | Donor 10 | Mean |
|---|---|---|---|---|---|---|---|---|---|---|---|
| QiaC | 7.04% | 7.47% | 5.00% | 7.93% | 7.81% | 4.91% | 11.80% | 5.82% | 8.39% | 10.67% | 7.68% |
| QiaM | 16.96% | 17.77% | 23.29% | 11.44% | 10.53% | 13.15% | 25.79% | 11.63% | 22.85% | 32.18% | 18.56% |
| SPRI | 17.63% | 15.98% | 21.59% | 18.70% | 11.44% | 12.55% | 24.51% | 13.64% | 22.68% | 37.10% | 19.58% |

### Supplemental Table 4 | Numerical values for Figure 2A and B.

| | | QIAM | | | | SPRI | | | | QIAM | | | | SPRI | | | |
|---|---|---|---|---|---|---|---|---|---|---|---|---|---|---|---|---|---|
| Donor | | DsDNase + | S1+ | Exo1 | Control | DsDNase + | S1+ | Exo1 | Control | Heatshock | No Heat Shock | dsDNA Library | Repair Enzyme | Heatshock | No Heat Shock | dsDNA Library | Repair Enzyme |
| 1A | uscfDNA (%integrated area of the intensity of 180-250bp) | 66.01 | 3.43 | 6.90 | 9.31 | 64.37 | 7.13 | 6.55 | 12.53 | 9.31 | 14.14 | 0.60 | 9.87 | 12.53 | 21.23 | 2.03 | 11.98 |
| | mncfDNA (%integrated area of the intenstiy of 250-400bp) | 21.38 | 56.79 | 51.50 | 49.2 | 29.23 | 47.70 | 43.49 | 43.44 | 49.20 | 37.39 | 37.20 | 46.57 | 43.44 | 12.37 | 41.98 | 40.41 |
| | uscfDNA fold signal from control or heatshock | 7.09 | 0.37 | 0.74 | 1.00 | 5.14 | 0.57 | 0.52 | 1.00 | 1.00 | 1.52 | 0.06 | 1.06 | 1.00 | 1.69 | 0.16 | 0.96 |
| | mncfDNA fold signal from control or heatshock | 0.43 | 1.15 | 1.05 | 1.00 | 0.67 | 1.10 | 1.00 | 1.00 | 1.00 | 0.76 | 0.76 | 0.95 | 1.00 | 0.28 | 0.97 | 0.93 |
| 2A | uscfDNA (%integrated area of the intensity of 180-250bp) | 69.13 | 3.53 | 5.87 | 11.93 | 48.15 | 7.16 | 4.99 | 11.12 | 11.93 | 17.96 | 1.85 | 8.01 | 9.12 | 26.28 | 2.88 | 8.01 |
| | mncfDNA (%integrated area of the intenstiy of 250-400bp) | 18.02 | 68.28 | 56.93 | 52.58 | 36.94 | 64.80 | 61.32 | 57.75 | 52.58 | 32.03 | 51.89 | 54.18 | 61.02 | 23.21 | 51.84 | 54.18 |
| | uscfDNA fold signal from control or heatshock | 5.79 | 0.30 | 0.49 | 1.00 | 4.33 | 0.64 | 0.45 | 1.00 | 1.00 | 1.51 | 0.16 | 0.67 | 1.00 | 2.88 | 0.32 | 0.88 |
| | mncfDNA fold signal from control or heatshock | 0.34 | 1.30 | 1.08 | 1.00 | 0.64 | 1.12 | 1.06 | 1.00 | 1.00 | 0.61 | 0.99 | 1.03 | 1.00 | 0.38 | 0.85 | 0.89 |
| 3A | uscfDNA (%integrated area of the intensity of 180-250bp) | 74.19 | 8.78 | 6.98 | 13.62 | 41.43 | 7.94 | 3.47 | 18.92 | 13.62 | 30.92 | 1.73 | 12.96 | 13.62 | 30.92 | 3.82 | 12.96 |
| | mncfDNA (%integrated area of the intenstiy of 250-400bp) | 14.43 | 61.55 | 59.99 | 54.73 | 45.39 | 55.38 | 57.62 | 52.9 | 54.73 | 31.80 | 45.66 | 50.99 | 54.73 | 31.80 | 53.96 | 50.99 |
| | uscfDNA fold signal from control or heatshock | 5.45 | 0.64 | 0.51 | 1.00 | 2.19 | 0.42 | 0.18 | 1.00 | 1.00 | 2.27 | 0.13 | 0.95 | 1.00 | 2.27 | 0.28 | 0.95 |
| | mncfDNA fold signal from control or heatshock | 0.26 | 1.12 | 1.10 | 1.00 | 0.86 | 1.05 | 1.09 | 1.00 | 1.00 | 0.58 | 0.83 | 0.93 | 1.00 | 0.58 | 0.99 | 0.93 |
| Mean | uscfDNA fold signal from control | 6.11 | 0.44 | 0.58 | 1 | 3.89 | 0.54 | 0.38 | 1 | 1.00 | 1.76 | 0.12 | 0.89 | 1.00 | 2.28 | 0.25 | 0.93 |
| | mncfDNA fold signal from control | 0.35 | 1.19 | 1.08 | 1 | 0.72 | 1.09 | 1.05 | 1 | 1.00 | 1.76 | 0.12 | 0.89 | 1.00 | 2.28 | 0.25 | 0.93 |
